# Distinct metabolite profiles in tissues and exudates of a monocot and dicot shaped by the environment

**DOI:** 10.1101/2024.08.30.610448

**Authors:** Alexandra Siffert, Sarah McLaughlin, Joëlle Sasse

**Affiliations:** Institute of Plant and Microbial Biology, University of Zurich, Zollikerstrasse 107, 8008 Zurich, CH

## Abstract

Plants exhibit remarkable plasticity in response to environmental changes. Understanding how plants adapt to diverse environmental conditions through changes in their metabolite profiles can provide insights into their adaptive strategies under suboptimal climate conditions. For this, metabolite profiles of tissues and root-derived, exuded compounds in various environmental conditions need to be characterized. Here, we compare the shoot, root, and root exudate metabolite profiles of the monocot *Brachypodium distachyon* and the dicot *Arabidopsis thaliana* grown in sterile, non-sterile, and sucrose-supplied basal salt medium or soil extract to represent natural and various standard laboratory conditions. We report unique metabolite fingerprints in shoots and roots for each species and environmental condition. Exuded compounds of Arabidopsis displayed higher sensitivity to soil extract conditions, whereas Brachypodium showed significant changes in response to non-sterile conditions. Organic acids, lipids, organic oxygen compounds, and phenylpropanoids were major contributors to the observed differences. Our results highlight the importance of considering environmental aspects when investigating plant metabolism and point towards crucial chemical classes involved in plant-microbe-environment interactions.

## 1 Introduction

Plants are sessile organisms, interacting with their environment by adjusting gene expression and metabolism, resulting in adaptation to e.g. nutrient availability or presence of microbe (Balmer *et al*., 2013; Erb and Kliebenstein, 2020; Sasse *et al*., 2018). The metabolism of different plant tissues has distinct functions in these interactions: leaves harbor photosynthesis, producing reduced carbon and energy equivalents that are further distributed to sink organs. Also, leaves defend against pathogens and herbivores by the production of secondary metabolites that function as deterrents and signals to other plant organs or adjacent individuals (Chen *et al*., 2020; Erb and Kliebenstein, 2020). Roots take up water and nutrients, defend against pathogens, and deliver reduced carbon to beneficial microorganisms either via specialized organs (root nodules, mycorrhizal interface) or by compounds into the soil surrounding roots, termed the rhizosphere. Further, roots export metabolites, so-called exudates. Exudates comprise a complex mixture of primary and secondary metabolites that are nutrients and signals to the root-associated community of organisms (Divekar *et al*., 2022; Rojas *et al*., 2014; Sasse *et al*., 2018; Wang *et al*., 2021b). Primary and secondary metabolites respond distinctly to different abiotic and biotic stimuli (Erb and Kliebenstein, 2020; Jan *et al*., 2021; Kumudini *et al*., 2018). For example, light response in leaves elevates glucose, fructose, and sucrose levels (Proietti *et al*., 2023), organic acids such as malate and citrate fluctuate diurnally (Fernie and Martinoia, 2009), and the amino acids glutamine and asparagine increase with abundant nitrogen (Yoneyama and Suzuki, 2020). Metabolite levels in root exudates also respond dynamically to nutrient levels and other abiotic stimuli (Salem *et al*., 2022). For example, elevated CO_2_ concentrations increased carbon exudation and soil organic matter turnover (van Groenigen *et al*., 2017), and the presence of *Bacillus subtilis* increased secretion of specific organic acids and phenolics that attract beneficial microbes resulting in enhanced plant growth (Badri *et al*., 2009), and *Lotus japonicus* growing in symbiosis with mycorrhiza stimulated exudation of arachidonic acids into the rhizosphere, attracting a microbiome beneficial for nutrient absorption (Xu *et al*., 2023).

In contrast to ubiquitously present primary metabolites, secondary metabolites are often produced only in response to stress conditions (Jan *et al*., 2021). They are central to plant defense, signaling, and interactions with the environment. For example, phenolic compounds such as flavonoids are produced in response to herbivory or pathogen attacks (Berhow and Vaughn, 1999; Kumar *et al*., 2023), and terpenoids such as limonene or pinene accumulate in leaves in response to physical damage to deter insects from feeding and to attract insect predators (Kegge and Pierik, 2010). Secondary metabolite production is often family-specific. For example, glucosinolates are powerful deterrents produced by the Brassicaceae family (Essoh *et al*., 2020), and legumes produce flavonoids that attract rhizobia (Abdel-Lateif *et al*., 2012). Thus, (secondary) metabolite profiles generally differ between plant species, ecotypes, and cultivars: Exudate profiles of grasses and forbs were distinct (Dietz *et al*., 2020), as were the exudation profiles of 19 *Arabidopsis thaliana* accessions (Mönchgesang *et al*., 2016). For 10 wheat genotypes, exudation was distinct in sugars, organic and amino acids, and in polyalcohols (Iannucci *et al*., 2017). Tissue and exudate metabolite profiles of *Brachypodium distachyon, Arabidopsis thaliana*, and *Medicago truncatula* shared approximately 60% of compounds, termed the core metabolome (McLaughlin *et al*., 2023b).

Although it is evident that metabolite profiles are distinct between plants and that environmental factors influence these profiles, few systematic studies have been done elucidating the importance of different chemical classes and metabolites. Thus, here we systemically compared shoot, root, and exudate metabolite profiles of two plant species across five growth environments. As growth environments, we chose typical laboratory conditions based on a basalt salt medium either in sterile or non-sterile conditions or supplemented with sucrose. As a more natural condition, we included sterile and non-sterile soil extract. We further investigated tissue and exudate profiles of *Arabidopsis thaliana* Col-0 lines established in different laboratories compared to two other ecotypes, La-0 and Wil1. We hypothesize that we see distinct metabolite profiles i) between tissues and exudates, ii) between plant species, iii) between growth conditions, iv) between Col-0 ecotypes, and v) similar profiles between different Col-0 laboratory lines.

## 2 Brachypodium and Arabidopsis tissue metabolite profiles are distinct and change specifically with environmental conditions

*Brachypodium distachyon* was chosen as a monocot representative and *Arabidopsis thaliana* as a dicot representative (here: Brachypodium, Arabidopsis). These two plant species are phylogenetically distinct and were shown to feature distinct metabolite profiles (McLaughlin *et al*., 2023b). They were grown for three weeks in a semi-hydroponic setup (McLaughlin *et al*., 2023a) in (a) ½ MS, (b) ½ MS non-sterile, (c) ½ MS + 1% sucrose, (d) 25% soil extract, and (e) 25% soil extract non-sterile. ½ MS is a minimal salt medium commonly used in controlled experimental conditions (Murashige and Skoog, 1962). Soil extract (SE) was prepared by incubating soil with water, followed by filter sterilization, resulting in a more natural but sterile environment comprising water-soluble soil compounds. Sucrose-supplemented conditions were included as a growth condition often used in laboratories (Badri *et al*., 2010; Chaparro *et al*., 2013; Micallef *et al*., 2009), and non-sterile samples were inoculated with a soil microbial extract to mimic a natural environment. Tissues were flash-frozen and extracted for metabolite analysis, and exudates were collected in ammonium acetate for 2 h.

Overall, 810-981 metabolites were detected in tissues, and 407-509 metabolites in exudates (Figure 1 a-f). The tissue metabolite profiles clustered separately for each species in a principal component analysis, as published previously (PCA, Suppl. Figure S.1 a-d) (Ayers and Thornton, 1968; Lesuffleur *et al*., 2007; Li *et al*., 2013; McLaughlin *et al*., 2023b; Micallef *et al*., 2009; Phillips *et al*., 2004; Strehmel *et al*., 2016; Sun *et al*., 2021). Interestingly, in Brachypodium and Arabidopsis, root and shoot SE samples clustered apart from ½ MS samples, demonstrating that the environment altered tissue metabolite profiles consistently across the two plant species investigated (Figure 1 a, b, d, and e). The sterile, nonsterile, and sucrose-supplemented conditions separated also, but less distinctly than ½ MS and SE (Figure 1 a, b, and d). Arabidopsis root ½ MS samples showed high variability and were thus excluded from the main PCA plot to showcase the distinctiveness of the other samples (Figure 1 e and Suppl. Figure S.1 i). To investigate whether metabolite profiles of SE-grown plants were similar to soil-grown plants, we included shoot metabolite profiles of pot-grown Brachypodium and Arabidopsis plants. Surprisingly, shoot metabolite profiles of soil-grown plants did not overlap with shoot profiles of SE-grown plants, a result consistent across both species (Suppl. Figure S.1 g and h). This suggests that other features aside from water-soluble compounds and microbial extracts impact shoot metabolite profiles.

**Figure 1:**
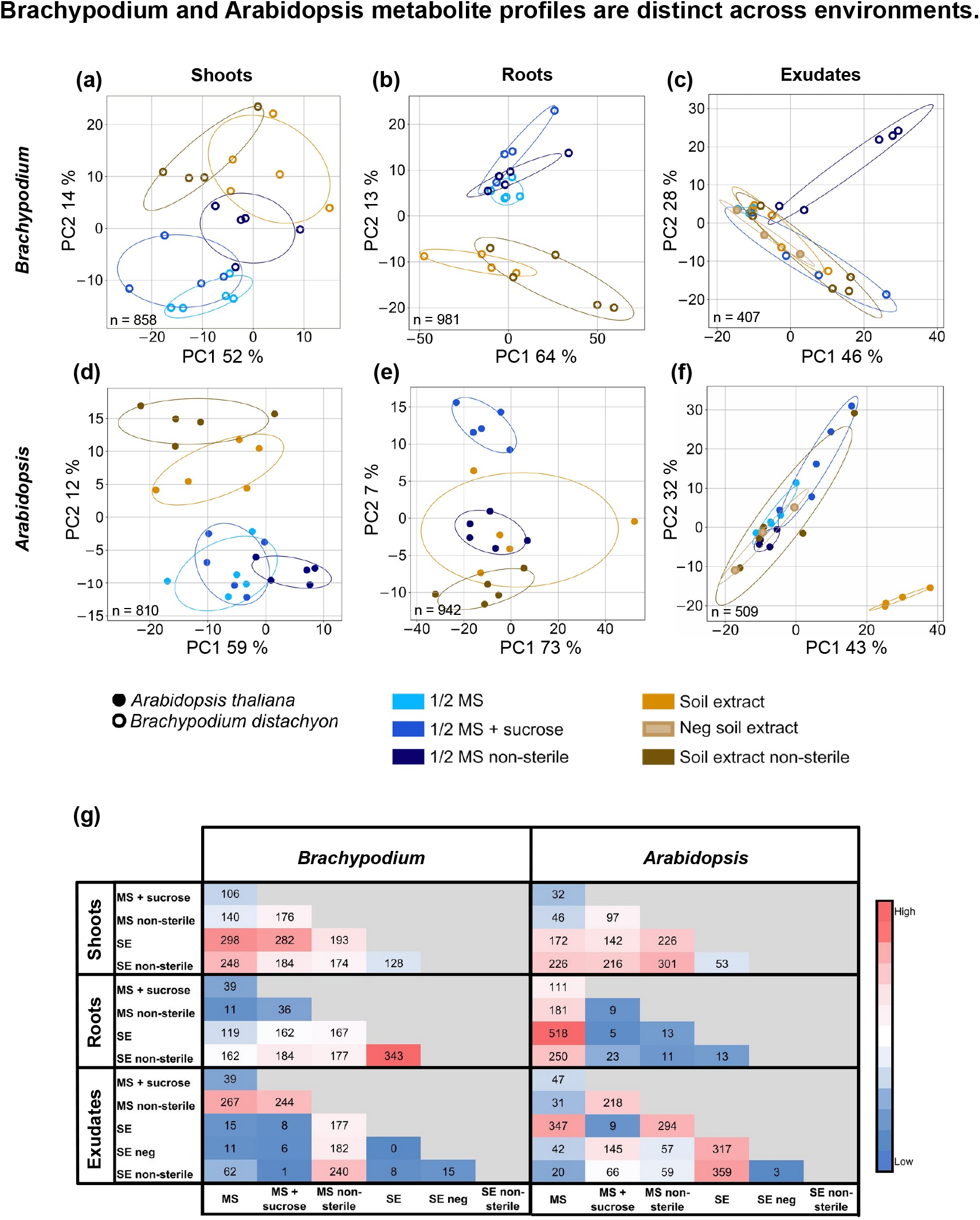
Brachypodium and Arabidopsis metabolite profiles are distinct across environments. (a-f) Principal component analysis (PCA) of metabolite profiles of shoots (a, d), roots (b, e), and exudates (c, f) of Brachypodium (a-c, empty circles) and Arabidopsis (d-f, full circles). Growth environments are indicated: ½ MS (light blue), ½ MS + sucrose (royal blue), ½ MS non-sterile (navy blue), SE (orange), SE non-sterile (brown), SE negative without plants (light brown). Number of metabolites for the analyses is indicated on respective graphs (n). Variances of the PCA are expressed in percent in Principal component (PC) 1 and PC 2. PCA plots including shoot profiles of soil-grown plants are presented in Figure S.1 g and h) Number of metabolites significantly changing between all pairwise comparisons in shoots, roots, and exudates of Brachypodium and Arabidopsis. Color code: grey: absent, red: high, blue: low. For percentages of metabolites significantly changing in comparison to ½ MS, see Suppl. Figure S.2. Data is based on 4-5 biological replicates (jars with each 3 Brachypodium or 5 Arabidopsis individuals).

In summary, our data confirmed species-specific tissue metabolite profiles. Further, we showed that tissue metabolite profiles of both species were responsive to all growth environments and that metabolite profiles of plants grown with hydroponic soil extract do not mirror profiles of plants grown on soil.

In our experimental setup, exudate profiles were less distinct between species compared to tissues (Suppl. Figure S.1 e and f). However, they were partially distinct when compared between different growth environments. Brachypodium ½ MS non-sterile and Arabidopsis SE exudate profiles separated the clearest from the rest of the samples (Figure 1 c and f and Suppl. Figure S.1 e and f). As soil extract consists of many metabolites, levels of exuded compounds need to be higher than in ½ MS to be detected against the soil metabolite background. For Arabidopsis, the only samples clearly clustering apart from negative control samples without plants were sterile SE conditions (no plants present, Suppl. Figure S.1 f). This data shows that changes in exudate metabolite profiles depend on the environmental conditions and are species-specific.

## 3 Natural versus laboratory conditions induce distinct metabolite profiles

We further investigated how many metabolites changed significantly between ½ MS and other environmental conditions (Figure 1 g). For Brachypodium, 9-26% of shoot, 1-14% of root, and 1-23% of exudate metabolites were distinct (Suppl. Figure S.2 a). For Arabidopsis, 3-20% of shoot, 10-45% of root, and 2-30% of exudate metabolites were distinct (Suppl. Figure S.2 a). The largest difference within the dataset was found between ½ MS vs SE conditions, specifically in shoots of Brachypodium and roots of Arabidopsis (26% and 45% of compounds, respectively, Suppl. Figure S.2 a). Shoots of both species featured similar changes in response to the growth environments, with a high number of metabolites changing between all pairwise comparisons of growth media, especially involving SE conditions (Figure 1 g). Brachypodium roots were mostly distinct in SE pairwise comparisons, and Arabidopsis roots in ½ MS pairwise comparisons (Figure 1 g). For exudates, Arabidopsis showed large percentages of metabolites changing in pairwise comparisons, whereas Brachypodium has rather low numbers except for ½ MS non-sterile conditions. It is interesting to note that ½ MS non-sterile conditions affects Brachypodium exudate metabolite profiles most, whereas Arabidopsis is more responsive to SE conditions.

## 4 Chemical classification of tissue metabolites is conserved across species and growth environments, whereas exudates are variable

In a next step, the chemical classes of metabolites by attribution to a Classyfire chemical superclass (Djoumbou Feunang *et al*., 2016), and 10 classes were detected overall (Figure 2 a-c). Shoot and root tissues shared the same chemical class footprint: Most abundant were organic acids and derivatives (19.8-21.7%), followed by lipids (18.6-25.4%), organoheterocyclic and organic oxygen compounds (15.1-16.2%, 12.5-14.9%), phenylpropanoids (9-11.2%), benzenoids (8.7-10.2%), nucleosides/nucleotides and analogs (3.6-4.5%), organic nitrogen compounds (1.4-1.8%), alkaloids, and other metabolites (1.1-1.3%) (Figure 2 c and Suppl. Figure S.3 a-d). The tissue footprints of the two species were not only identical across tissues, but were also conserved in the different growth environments.

**Figure 2:**
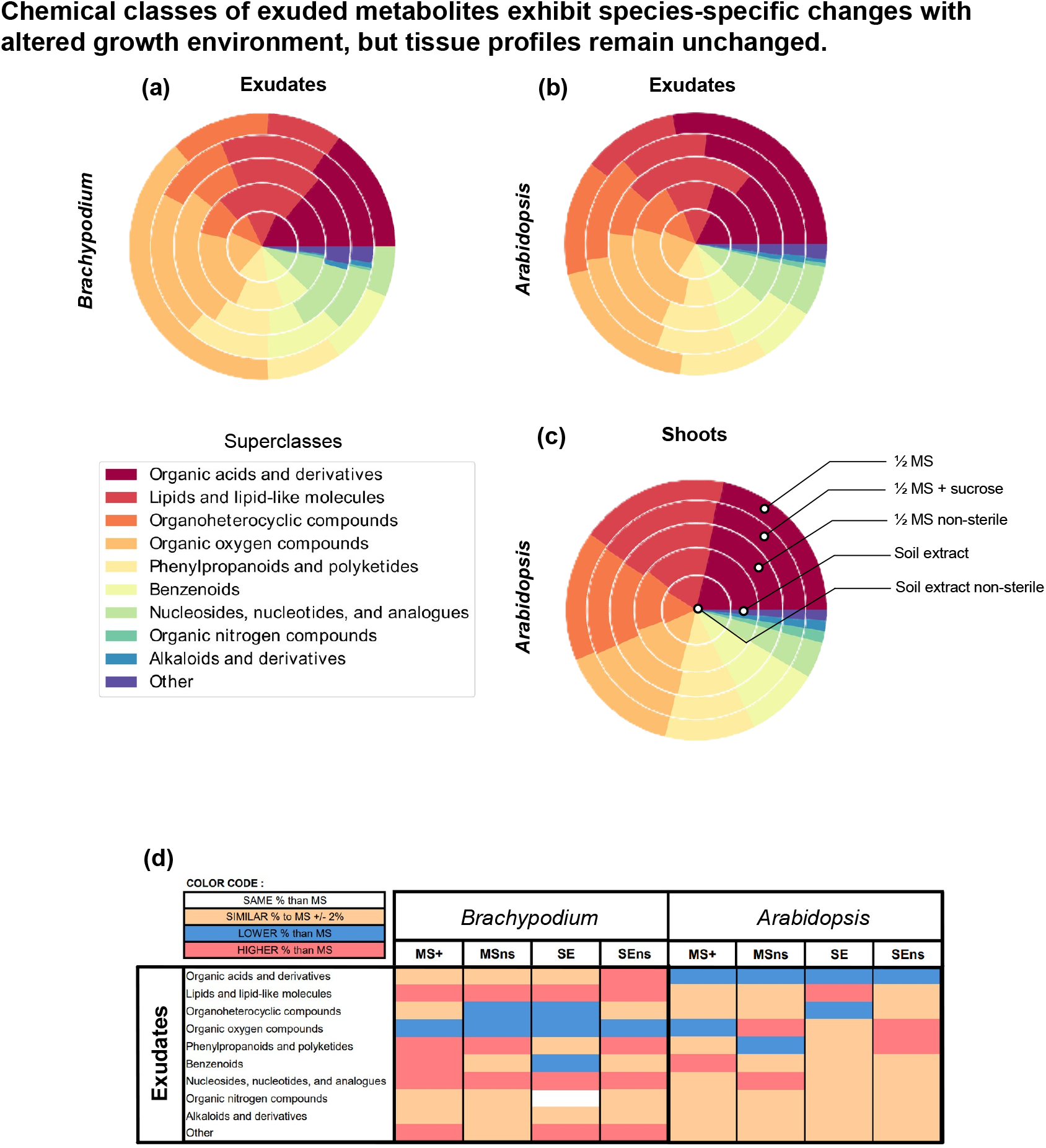
Chemical classes of exuded metabolites exhibit species-specific changes with altered growth environment, but tissue profiles remain unchanged. (a-c) Pie charts of chemical superclasses of exudates (a, b) and tissues (c) of Brachypodium (a) and Arabidopsis (b, c) (Classyfire: https://cfb.fiehnlab.ucdavis.edu/, Djoumbou Feunang *et al*. (2016)) based on metabolite presence in the different samples. The five circles represent the different growth environments, from outside to inside: ½ MS, ½ MS + sucrose, ½ MS non-sterile, SE, SE non-sterile. The chemical classes are indicated by color. Other classes comprise non-identified compounds, organophosphorus compounds, hydrocarbon derivates, lignans, and neolignans and related compounds. Additional tissue chemical class plots are presented in Suppl. Figure S.3 b, c, and d. See Figure 1 for PCA analyses of data. See Suppl. Figure S.3 for comparison of Brachypodium and Arabidopsis tissues. (d) Comparison of chemical class abundance in percent in various environments versus ½ MS in Brachypodium and Arabidopsis exudates. Color code: red: chemical class more abundant compared to ½ MS (>2% difference), blue: chemical class less abundant (>2% difference), orange: chemical class as abundant (<2% difference), white: chemical class the same as in ½ MS. See Suppl. Figure S.3 e for number of metabolites assigned to the different chemical classes.

In exudates, the same chemical classes were detected as in tissues. In contrast to the robust tissue footprint however, relative proportions of exuded metabolites were distinct from tissues and further also differed between growth environments (Figure 2 a, b, and d). Nucleosides/nucleotides and lipids increased and organoheterocyclic compounds decreased in several environmental conditions compared to ½ MS across both species. Further, organic nitrogen and alkaloid compounds did not change in either species. Aside from these findings, no responses were shared: In Brachypodium, lipid and nucleoside exudation increased and organic oxygen compounds decreased when comparing ½ MS to other environments (Figure 2 a and d). In Arabidopsis, organic acid exudation decreased in all growth environments compared to ½ MS (Figure 2 b and d). Other chemical classes showed a more mixed response: Sucrose addition to ½ MS resulted in less organic oxygen and more benzenoid exudation, and growth in SE resulted in elevated lipid and reduced organoheterocyclic compound exudation. Non-sterile growth conditions elevated organic oxygen exudation but had contrasting effects on phenylpropanoid exudation: in non-sterile ½ MS, phenylpropanoid exudation was reduced whereas in non-sterile SE, their exudation was increased. (Figure 2 d). Interestingly, Arabidopsis exudates of non-sterile SE conditions were comparable to Brachypodium exudates of non-sterile MS, SE, and ½ MS + sucrose conditions, featuring increased phenylpropanoid exudation (Figure 2 d).

When comparing changes in chemical classes versus overall changes in metabolite profiles, different trends are observed (Figures 1 and 2): the number of chemical classes changing is not a direct indicator of the distinctiveness of metabolite profiles, nor does a distinct metabolite profile translate to a distinctiveness in chemical classes. Although tissue metabolite profiles were distinct between species, tissues, and environmental conditions, their chemical class footprint was conserved. In contrast, chemical classes differed between exudates of both species and environmental conditions, whereas their metabolite profiles were similar in a PCA analysis. We conclude that for a thorough analysis, both, full metabolite profiles as well as chemical classification should be analyzed.

## 5 Organic acids, lipids, organic oxygen compounds, and phenylpropanoids drive distinct exudate profiles associated with growth environments

We further investigated which metabolites were central to the changes in metabolite profiles observed. For this, we compared exudate (Figure 3 a and b) and tissue (Suppl. Figure S.4 a-d) profiles of ½ MS versus the other environments. As a general trend, abundance of many metabolites increased in exudates of both species grown in environments other than ½ MS (Figure 3).

**Figure 3:**
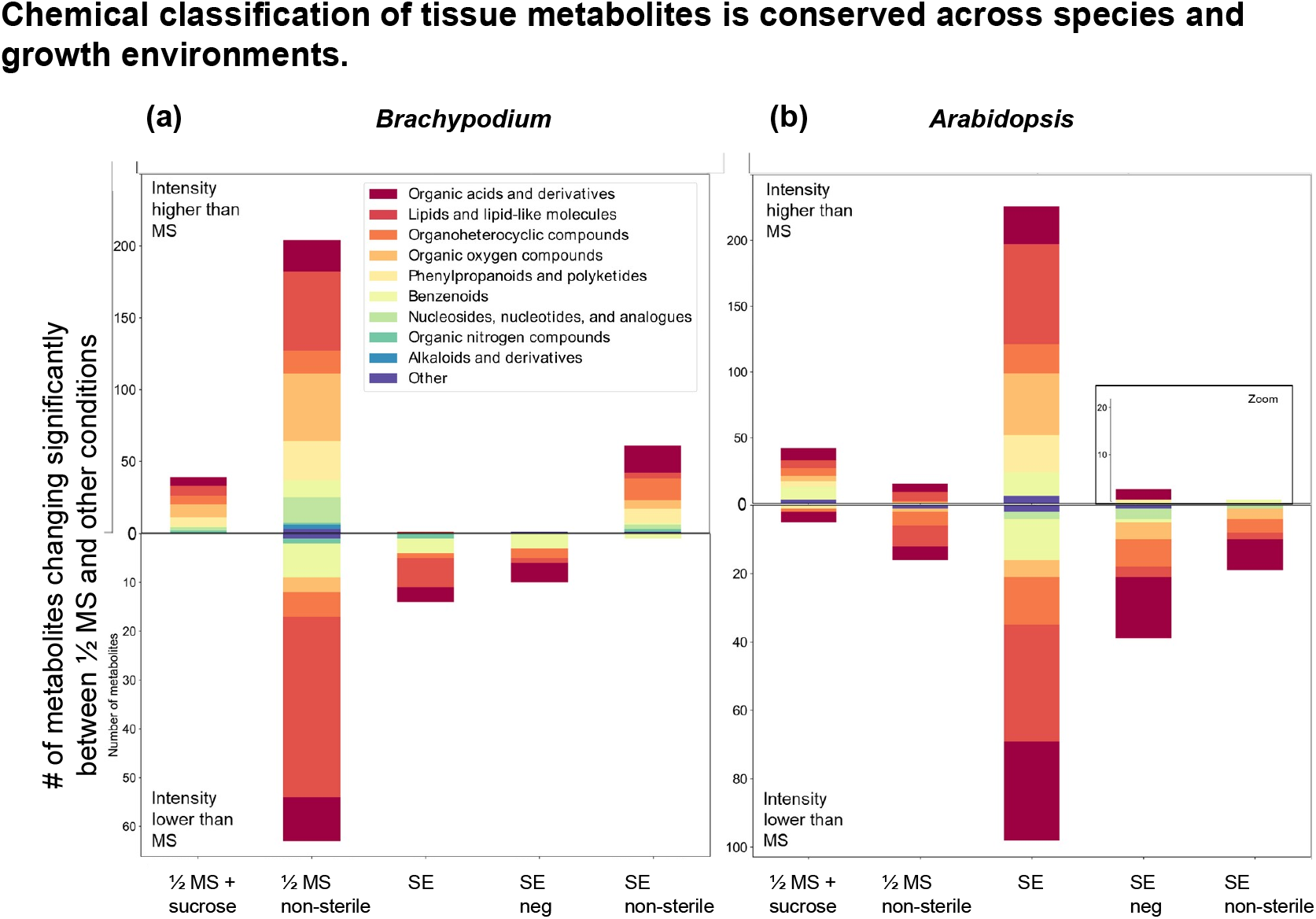
Chemical classification of tissue metabolites is conserved across species and growth environments. (a-b) Number of metabolites changing significantly between ½ MS and other environments for Brachypodium (a) and Arabidopsis (b) root exudates. Number of more and less abundant compounds are indicated on positive and negative axes, and metabolites are colored according to superclass (see also Figure 2, Suppl. Figure S.3, a and b for amount of metabolites significantly changing). For metabolites changing in tissues, see Suppl. Figure S.4. For number of changing metabolites see Suppl. Figure S.6 a and b.

In Brachypodium exudates, most compounds changed in ½ MS versus ½ MS non-sterile conditions (see also Figures 1 and 2), with 204 compounds across all 10 chemical classes being more abundant in nonsterile conditions (76.4%, total 267 compounds, Figure 2 g, Figure 3 a). Phenylpropanoids, nucleosides, and alkaloids increased in abundance, whereas SE and SE negative controls featured mostly compounds with reduced abundance. Organoheterocyclic, organic oxygen, and nucleoside compounds also generally increased in abundance. Other chemical classes showed distinct responses: organic acid exudation decreased in SE, and increased in ½ MS + sucrose and SE non-sterile conditions (Suppl. Figure S.6 a). 55 lipids increased in ½ MS non-sterile conditions, and 37 decreased in abundance.

In Arabidopsis exudates, most compounds changed in SE versus ½ MS (see also Figures 1 and 2). Of these, 248 compounds increased, and 99 compounds decreased in abundance, and changes observed across all 10 chemical classes (71% and 29%, respectively Figure 3 b). SE non-sterile samples only featured metabolites with increased abundance, comprising organic acids, organoheterocyclic, and organic oxygen compounds (Suppl. Figure S.6 b). Organic acids and heterocyclic compounds were more prominent contributors to change in Arabidopsis compared to Brachypodium, and lipids contributed to change equally in both species (Figure 3).

In shoots, many distinct compounds were detected, showing a mirrored pattern of compounds increasing and decreasing across the environmental conditions (Suppl Figure S.4 a and b and Suppl. Figure S.5 a). Brachypodium featured many distinct compounds in SE compared to ½ MS (298 and 248 metabolites for SE and SE non-sterile, Suppl. Figure S.5 a). Similarly, Arabidopsis also featured many distinct compounds in SE conditions compared to ½ MS (172 and 226 metabolites, respectively) (Suppl Figure S.4 e and f and Suppl. Figure S.5 d).

In contrast to shoots, root metabolite patterns are largely distinct between both species. Brachypodium featured a high number of distinct compounds in SE conditions compared to ½ MS, but few changes in the other conditions (Suppl. Figure S.4 c and d). Root metabolite abundance generally decreased in SE and increased in SE non-sterile, with trends being consistent across chemical classes. Arabidopsis roots somewhat mirrored the trend for SE decreased and SE non-sterile increased abundance, but less prominently (Suppl. Figure S.4 g and h).

Overall, organic acids, lipids, organic oxygens, and phenylpropanoids were the chemical classes that contributed most to significant changes in Arabidopsis and Brachypodium (Figure 3 a and b and Suppl. Figure S.6 a and b). In our dataset, organic acids were mostly composed of carboxylic acids, such as amino acids and derivatives. Lipids contained fatty acyls, prenols lipids (mostly terpenoids), and steroid derivates. Organic oxygen compounds contained mostly carbohydrates (monosaccharides and aminosaccharides), and phenylpropanoids mostly comprised of flavonoids and stilbenes (Suppl. Figure S.5). When comparing Brachypodium in ½ MS non-sterile and Arabidopsis SE conditions against ½ MS, nutrients such as carbohydrates largely increased in exudates (Suppl. Figure S.6 c). Of the 42 carbohydrates detected in Brachypodium and Arabidopsis, all of them are differentially released: 37 carbohydrates were shared between species, with 5 being specific to Arabidopsis.

Overall, 82-107 amino acids were detected in tissues, with similar numbers for Brachypodium and Arabidopsis (Figure 4 a). With our method, most single amino acids such as alanine, serine, proline, valine, cysteine, leucine, asparagine, histidine, phenylalanine, tyrosine, arginine, and glutamine were detected, and changes in abundance for single compounds are seen. For example, methionine levels significantly increased in Brachypodium shoots in ½ MS non-sterile conditions, and in Arabidopsis SE and SE nonsterile conditions (Suppl. Figure S.7 g and h). Exudate profiles comprised 0-64 amino acids, with fewer amino acids being detected in Brachypodium exudates compared to Arabidopsis (0-18 in Brachypodium, 14-64 in Arabidopsis, Figure 4 a, b). Only 9 amino acids were differentially exuded by both species, with an addition 16 amino acids specifically by Arabidopsis, and 1 by Brachypodium. Amino acids increased in Brachypodium ½ MS non-sterile samples, but mostly decreased in Arabidopsis SE samples.

**Figure 4:**
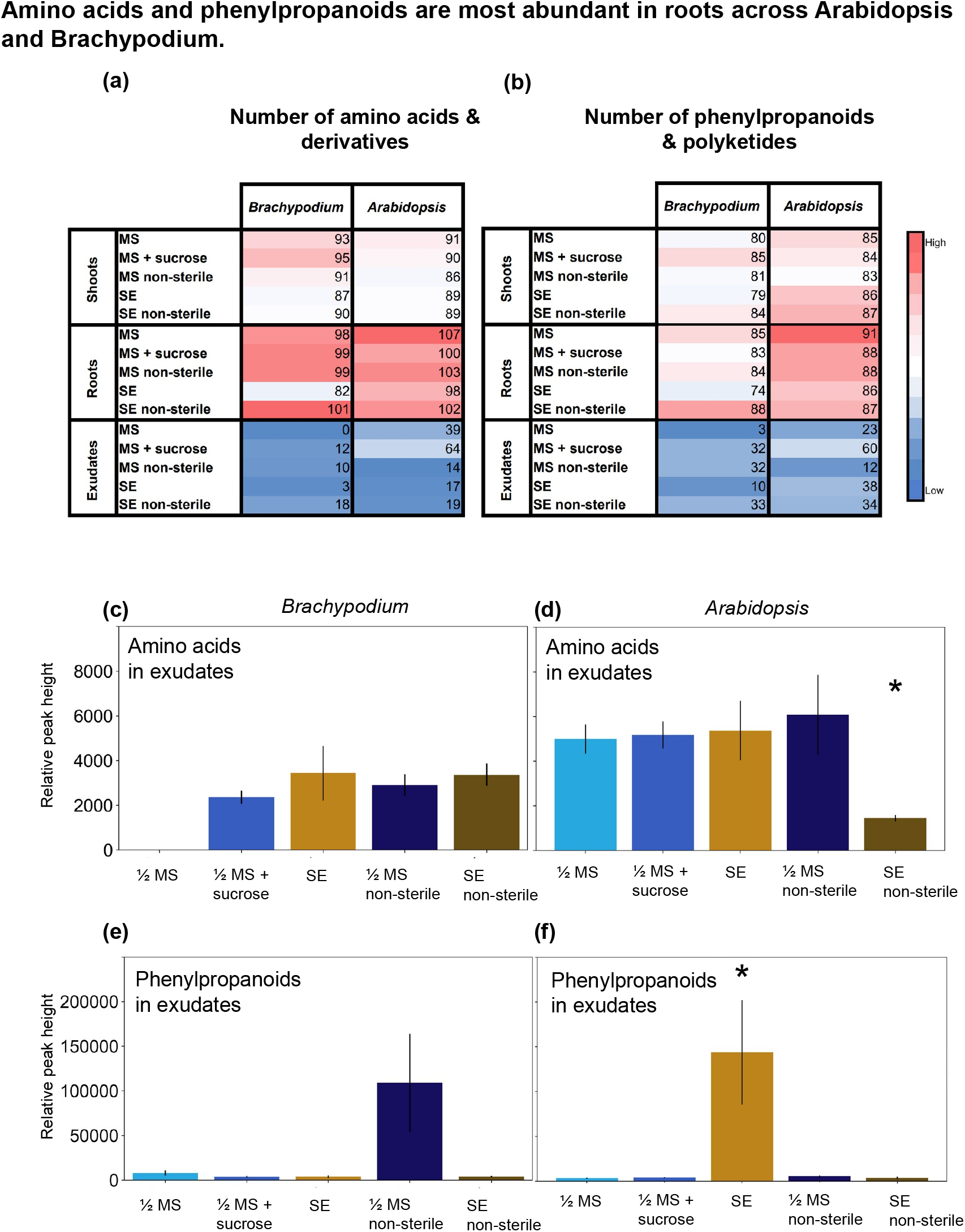
Amino acids and phenylpropanoids are most abundant in roots across Arabidopsis and Brachypodium. (a-b) Number of amino acids and derivatives (a) and phenylpropanoids and polyketides (b) detected in Brachypodium and Arabidopsis tissues and exudates grown in five distinct environments. Red colors indicate high, white intermediate, and blue low numbers. The color map is specific to each sample type. (c-f) Examples of exuded amino acid (c, d) and phenylpropanoid (d, f) compounds in exudates of Brachypodium (c, e) and Arabidopsis (d, f). Data is displayed as average ± S.E.M., *: pvalue < 0.05 (T-Test vs. ½ MS).

Phenylpropanoids were also more abundant in tissues than exudates: 74-91 compounds were detected in tissues of both species, and 3-60 compounds in exudates (Figure 4 b). Of the defense compounds detected, flavonoids, stilbenes, steroids and derivatives accumulated in exudates of both species (Brachypodium ½ MS non-sterile, Arabidopsis SE, Suppl. Figure S.6 c). Terpenoids showed a mixed response with two thirds of compounds increasing, and one third decreasing in abundance. Exudation of these compounds could be stimulated by microbes in nonsterile environments (Brachypodium ½ MS non-sterile), or by microbial elicitors (Arabidopsis SE).

The number of phenylpropanoids and amino acids was similar for tissues of both plant species. In contrast, numbers of these metabolites detected in exudates were specific to each environmental condition and species. Both, amino acid and phenylpropanoid exudation increased in ½ MS + sucrose compared to ½ MS condition for both species (Figure 4 a and b). Interestingly, although more amino acid compounds were exuded, amino acid levels only increased in Brachypodium exudates (Figure 4 c), and not in Arabidopsis exudates (Figure 4 d-f). Similarly, the increased phenylpropanoid compound numbers also did not result in elevated phenylpropanoid abundance. For the latter cases, this suggests that the increased number of compounds was compensated with reduced exudation levels, resulting in a similar total amount of compounds exuded.

## 6 Discussion

Our study investigated how plants respond on a metabolic level to different environmental conditions. We found that metabolite profiles of shoots and roots are distinct for each condition tested. Biggest differences are observed between ½ basal salt medium and soil extract conditions, caused by distinct compounds being present and by distinct metabolite abundances. Interestingly, the chemical class fingerprints remained the same across environments and plant species. Exudate metabolite profiles were more similar across environments and species. However, in contrast to tissue samples, the relative abundance of chemical classes in exudates changed across environments and plant species. Two conditions, ½ MS non-sterile for Brachypodium and SE for Arabidopsis, were most distinct for exudates from ½ MS sterile conditions. Thus, the microbes present in ½ MS non-sterile and the microbe-derived fragments in SE conditions might have triggered the similar metabolite changes in the two plant species investigated.

In exudates of both species, lipids, amino acids, and phenylpropanoids changed significantly. In Arabidopsis soil extracts (SE) conditions, despite no observable infection traits, defense compounds like flavonoids, stilbenes, and coumarins increased, suggesting that bacteria, whether beneficial or pathogenic, triggered an immune response. However, in non-sterile conditions harboring alive microbes, no release of defense compounds was observed for Arabidopsis. This might indicate a complex interplay between microbe-derived signals and alive bacteria, which could be interrogated in future experiments. Brachypodium featured an opposite pattern compared to Arabidopsis with defense compounds exuded in non-sterile conditions, but not in SE conditions. The amount of defense-inducing elicitors in soil microbial communities, in sterile soil extract, and their interplay with beneficial and/or neutral microbes remains to be investigated. To distinguish defense compound production in response to pathogens and other microbes, plants could be inoculated with single pathogenic or beneficial strains to investigate changes in exudation and tissue metabolite profiles, coupled with phenotypic responses and microbial growth. Further, to exclude microbial metabolism from the experiment and only investigate plant responses, plants could be inoculated with a microbial extract instead (Portieles *et al*., 2021). Further, extension of this study to additional monocots and dicots would provide information whether these differences observed are conserved or specific to the investigated species.

Phenylpropanoids, similar to other defense compounds, increase in response to microbial stimuli (Balmer *et al*., 2013; Kumar *et al*., 2023). They can act as toxins, for example discouraging herbivory feeding, or they feature antimicrobial and antifungal properties (Guerriero *et al*., 2018). Typical defense compounds of grasses are benzoxazinoids, which are synthesized in leaves, and transported throughout the plant, including roots and exudates (Niculaes *et al*., 2018). Arabidopsis synthesizes glucosinolates primarily in leaves, but also in roots, flowers, and seeds depending on the development stage (Kopriva and Gigolashvili, 2016). In our data, glucosinolates indeed increased in Arabidopsis exudates and roots in non-sterile and SE conditions.

We further detect a plethora of changes in presence and abundance of primary metabolites in the different growth environments. Changes in organic acids might relate to nutrient deficiencies: exudation of organic acids is often upregulated in response to phosphate deficiency, as carboxylates can solubilize phosphate bound to soil minerals (Vengavasi *et al*., 2021). In our dataset, increased organic acid exudation is observed in non-sterile conditions, which could point towards a competition with microbes for resources (Yamada and Osakabe, 2018). Amino acid exudation also increased. Some amino acids are osmolytes and osmoprotectants, their increased exudation could be a response to suboptimal osmotic conditions in the various environments (Dikilitas *et al*., 2020; Kebert *et al*., 2022). In nonsterile or SE environments, changes in plant metabolism might be a response to microbial presence. Exudation of carbohydrates and other reduced carbon compounds was increased when Brachypodium and Arabidopsis were in contact with bacteria, which might indicate that plants modulate exudation to attract beneficial microbes.

Future studies could expand the research presented here in several ways. On the technical side, different chromatography methods targeting different types of compounds should be employed to capture more of the metabolite diversity present, including polar and nonpolar molecules, gaseous and polymeric substances. To complement the picture of enzymes, proteins, RNA, and DNA (Wen *et al*., 2009) released from root-derived cells should be characterized in different environments to capture the full effect of plant-derived molecules changing in distinct environments. For this, it would be useful to build a comprehensive database for plant-derived compounds including ideally quantitative information comprising representative species grown in various settings.

Our study is a first step towards revealing distinct responses of plants grown in controlled and natural conditions, which is a central aspect when translating findings into applications. While laboratory settings allow for precise control and replication, they do not fully capture the complexity of interactions seen in natural soils, leading to distinct metabolic profiles. The observed differences in metabolite profiles between plants grown in pots versus natural soil conditions likely stem from variations in root architecture, microbial communities, and nutrient availability. These findings suggest that future studies should carefully consider experimental design, particularly the choice of growth medium, to accurately reflect the metabolic responses of plants in more naturalistic settings.

## Acknowledgments

We thank Prof. Dr. Nicola Zamboni from the ETH Zürich, Switzerland for determining the root exudation profiles with direct injection. We thank Dr. Yvonne Steinbach from the University of Zurich, Switzerland for discussions regarding experimental setups. Further, we acknowledge the Swiss National Science Foundation (PR00P3_185831 to J.S., supporting A.S. and S. M. and J.S.).

## 7 Materials and methods

### Plant material and growth media

*Arabidopsis thaliana* Col-0 and *Brachypodium distachyon* Bd21-3 were used in the experiments presented. For the Arabidopsis ecotype and laboratory lines experiment, Col-0 lines were collected from 5 laboratories at University of Zurich, 1 from ETH of Zurich, 1 from University of Basel, and 1 from Carnegie Institution for Science, CA, USA. La-0 and Wil-1 ecotypes were used as outgroups.

The growth medium was: ½ MS: half-strength Murashige and Skoog medium (MS, M0221.0050, Duchefa Biochemie, Haarlem, Netherlands), ½ MS + 1% sucrose (P15030B, AppliChem, Darmstadt, Germany). The soil extract SE was prepared by dissolving 800 g of sieved soil in 1 L of milliQ water for 16 h at 4°C and 300 rpm, followed by filtration through a filter paper (Whatman 1001-150, GE Healthcare Life Sciences, Milwaukee, Wisconsin, USA) and sterile filtration through a 0.22 µm filter (S2GPU11RE, Sigma Aldrich, Saint-Louis, USA). Plants were inoculated with 1:4 diluted SE in milliQ water. Non-sterile samples were prepared as follows: a soil slurry was prepared as described above. The 0.22 µfilter paper was incubated in 10 ml milliQ water and vortexed for 5 min. The resulting microbial slurry was inoculated 1:100 into the growth media.

### Plant growth conditions

Arabidopsis seeds were sterilized for 15 min in 70 % ethanol, removed to 100 % ethanol for 15 min, removed, and allowed to dry on a sterile bench for 3 h. Brachypodium seeds were sterilized for 30 sec in 70 % ethanol, rinsed 5 times with sterile MilliQ, then incubated for 5 min in 10 % bleach, and finally rinsed 5 times with MilliQ. Dry seeds were germinated on ½ MS plates and placed in growth chamber (Conviron 8/10, 16 h light/8 h dark, 22 °C day/18 °C night, 150160 µmol m^-2^ s^-1^).

Plants were germinated on ½ MS plates supplied with 0.7% phytoagar (half-strength Murashige and Skoog medium: M0221.0050, Duchefa Biochemie, Haarlem, Netherlands, phytoagar: P10003.1000, Duchefa Biochemie, Haarlem, Netherlands). Brachypodium plants were transferred at 5d and Arabidopsis at 15d into a semi-hydroponic setup with 35 ml growth medium (McLaughlin *et al*., 2023a). Plants were grown in a growth chamber (Conviron 8/10) under 16 h light and 8 h dark conditions, at 22 °C during the day (150-160 µmol m^-2^ s^-1^) and at 18 °C during the night.

### Experimental design

For each environmental condition, 5 biological replicates (jars) each consisting of 3-5 plants were prepared. Negative controls comprised 3 replicates (jars) to capture the metabolite background of the respective systems. The semi-hydroponic system was prepared in sterile conditions (McLaughlin *et al*., 2023a). Sterility was confirmed by plating a 20 µl aliquot on LB medium (Bactotryptone: 84610.0500, Avantor by VWR Chemical, Rosny-sous-Bois, France, NaCl: 3957.1, Carl Roth, Arlesheim, Switzerland, bacto-yeast extract: 84601.0500, Avantor by VWR chemical, Rosny-sous-Bois, France), incubating the plate for 48h at 23°C, and monitoring for microbial growth.

### Metabolite sample collection

The growth media was exchanged with 50 ml of filter-sterilized 20 mM Ammonium acetate pH 5.7 (102308585, Sigma-Aldrich, Missouri, USA) as described (McLaughlin *et al*., 2023a). Exudates were collected for 2 h. At 19-21 dag, the shoots and root fresh weight of plants were recorded, and 250 mg of tissues were frozen in liquid nitrogen for metabolite extraction. The extraction was performed on ice with 2x 800 ml of MeOH:ACN:H2O (4:4:2) for 1 hour with intermittent vortexing. The supernatant was dried via speed vac (UNIVAPO 100 ECH, UniEquip, Bayern, Germany). Tissue samples were dissolved in MEOH:H2O 1:1 with a concentration of 70 mg 200 µL^-1^ and diluted to 1:150. Root exudate volumes were normalized with 20mM Ammonium acetate according to total root mass.

Two technical replicates were injected into a flow injection time of flight mass spectrometry system of an isocratic Agilent 1200 pump coupled to a Gerstel MPS2 autosampler and an Agilent 6550 QTOF mass spectrometer (Agilent, California, USA). The platform operated with published settings (Fuhrer *et al*., 2011). The isocratic flow rate was 150 µl min^-1^ of mobile phase, consisting of 60:40% v/v isopropanol:water buffered with 5 mM ammonium fluoride for negative ionization mode. For online mass axis correction, homoserine and Hexakis (1H, 1H, 3H-tetrafluoropropoxy)-phosphazine were used. Mass spectra were recorded from 50-1000 m/z with a frequency of 1.4 spectra/s using the highest available resolving power. Source temperature was set to 325°C, with 5 L/min drying gas and a nebulizer pressure of 30 spig. Fragmentor, skimmer, and octupole voltages were set to 175 V, 65 V, and 750 V, respectively. Spectral data was processed and annotated as published based on the Kyoto Encyclopedia of Genes and Genomes (KEGG) and Human Metabolome Database (HMDB) databases (Fuhrer *et al*., 2011).

### Metabolite Data Analysis

Quality control samples were checked for m/z and intensity shifts. Blank samples were injected after quality control samples, and after each sample group (e.g. roots, shoots, exudates, different experiments) to avoid carryover. Data processing was completed in Visual code studio with Python. The two technical replicates for each sample were averaged (sample mean), and the mean of each species or negative control was calculated. Metabolites were deemed present in the dataset if the mean of the metabolite abundance in a single plant species was two times higher than the mean of negative control. The intensity of filtered metabolites underwent percentage normalization for PCA analysis. Statistical analysis (ANOVA and post hoc TUKEY tests) was completed for each metabolite present in the dataset. For the assignment of chemical classes to compounds, KEGG IDs were converted to InChIKeys, which was used as an input for Classyfire (https://cfb.fiehnlab.ucdavis.edu/, Djoumbou Feunang *et al*. (2016)). For the determination of metabolite presence or absence in the different plant species, metabolites were deemed to be present if their mean intensity in the species of interest was higher than two times the mean intensity. The following Python packages were used; pandas, numpy, scipy, matplotlib, sklearn, statsmodels, and multiprocessing.

## Supplementary material

**Figure S.1:**
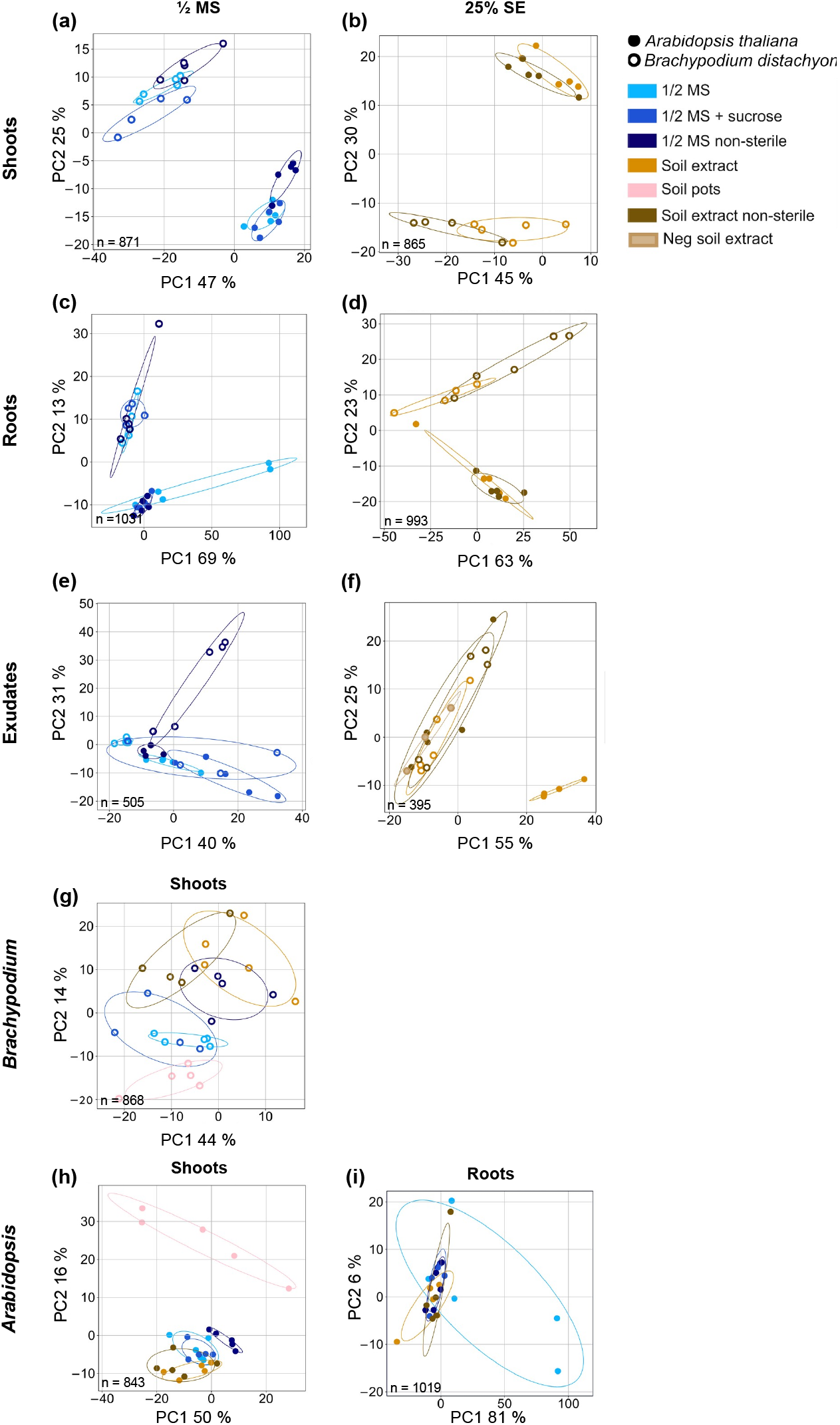
Brachypodium and Arabidopsis metabolite profiles are distinct across environments. (a-f) Principal component analysis (PCA) of metabolite profiles of shoots (a, b), roots (c, d), and exudates (e, f) of Brachypodium (empty circles) and Arabidopsis (full circles) separated according to environments conditions. Shoots metabolites of Brachypodium and Arabidopsis are displayed again with the addition of the soil-grown respective plants for comparison (g, h). The Arabidopsis root metabolite PCA which expresses high variability in Col-0 is provided in i. Growth environments are indicated: ½ MS (light blue), ½ MS + sucrose (royal blue), ½ MS non-sterile (navy blue), SE (orange), SE non-sterile (brown), SE negative without plants (light brown), soil-grown plants (pink). Number of metabolites for the analyses is indicated on respective graphs (n). Variances of the PCA are expressed in percent in Principal component (PC) 1 and PC 2.

**Figure S.2:**
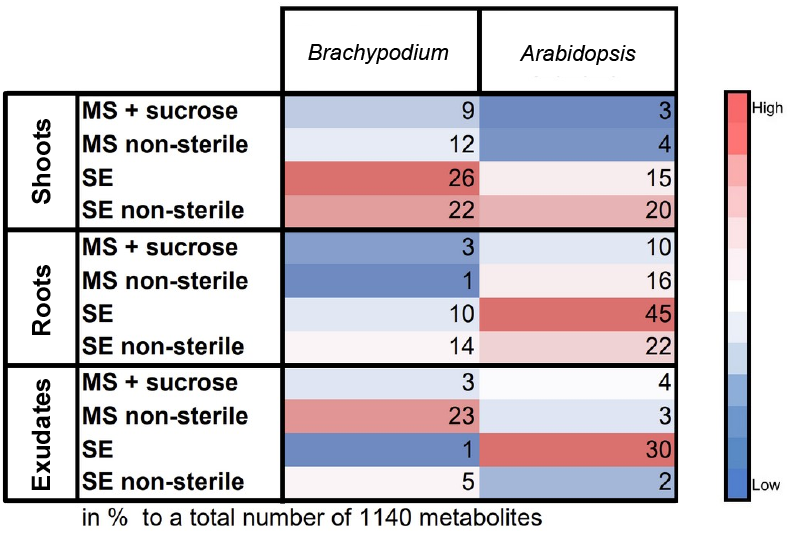
Significantly changing metabolites in comparison to ½ MS. Percentages of metabolites significantly changing in comparison to ½ MS for shoots, roots, and soil exudates of Brachypodium (left) and Arabidopsis (right). Color map based on tissues and exudates. The percentage is based on a total number of metabolites of 1140. Color code: red: high, blue: low.

**Figure S.3:**
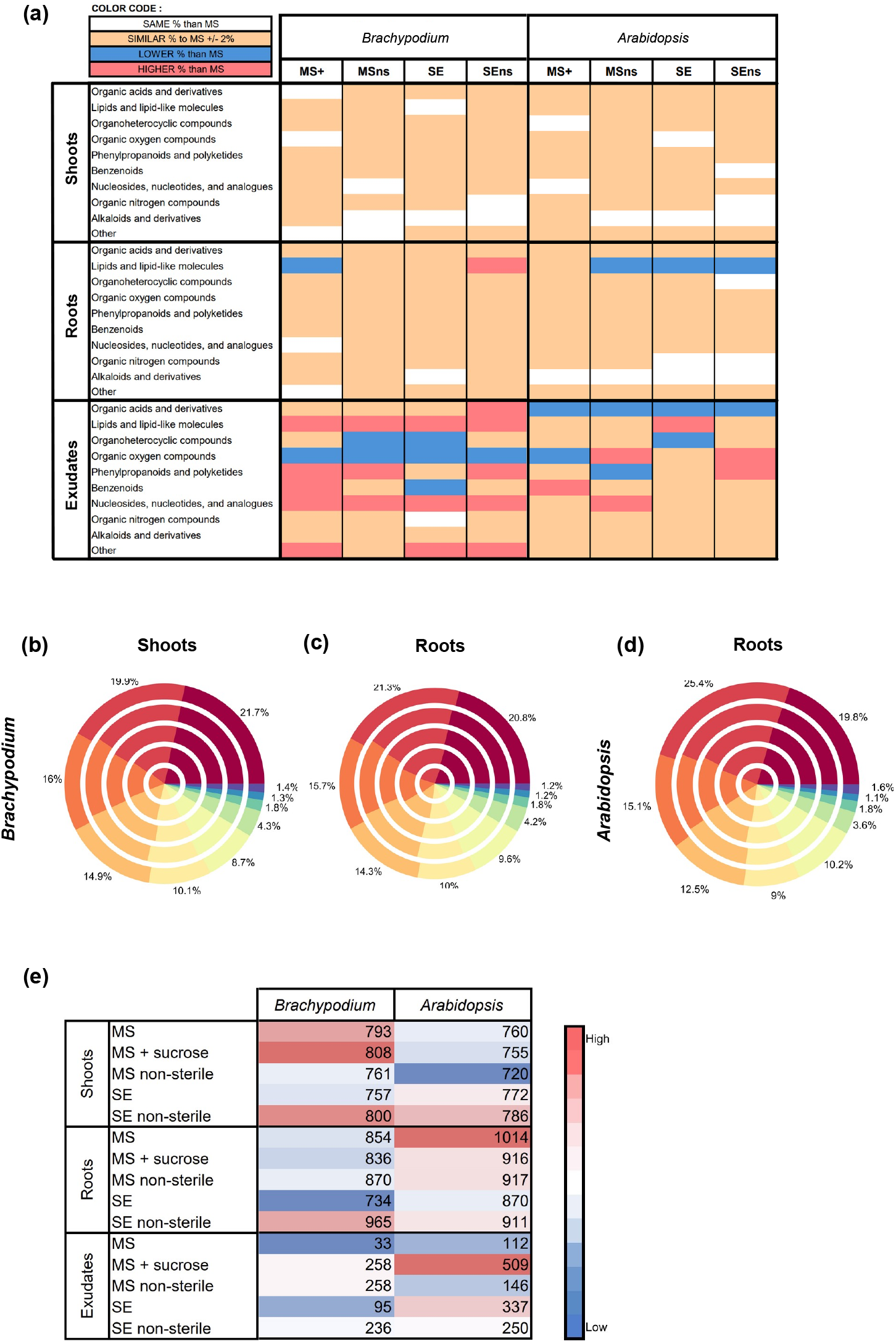
Pie charts of chemical superclasses of tissues and exudates from Brachypodium and Arabidopsis grown in 5 distinct environments and comparisons to ½ MS conditions. (a) Comparison of chemical class abundance in percent in various environments versus ½ MS in Brachypodium and Arabidopsis exudates. Color code: red: chemical class more abundant compared to ½ MS (>2% difference), blue: chemical class less abundant (>2% difference), orange: chemical class as abundant (<2% difference), white: chemical class the same as in ½ MS. (b-c) Pie charts of chemical superclasses of and tissues of Brachypodium (b, c) and Arabidopsis (d) (Classyfire: https://cfb.fiehnlab.ucdavis.edu/, Djoumbou Feunang *et al*. (2016)) based on metabolite presence in the different samples. The five circles represent the different growth environments, from outside to inside: ½ MS, ½ MS + sucrose, ½ MS non-sterile, SE, SE non-sterile. The chemical classes are indicated by color. Other classes comprise non-identified compounds, organophosphorus compounds, hydrocarbon derivates, lignans, and neolignans and related compounds. (e) Number of metabolites present for each environmental condition used to build all the pie charts.

**Figure S.4:**
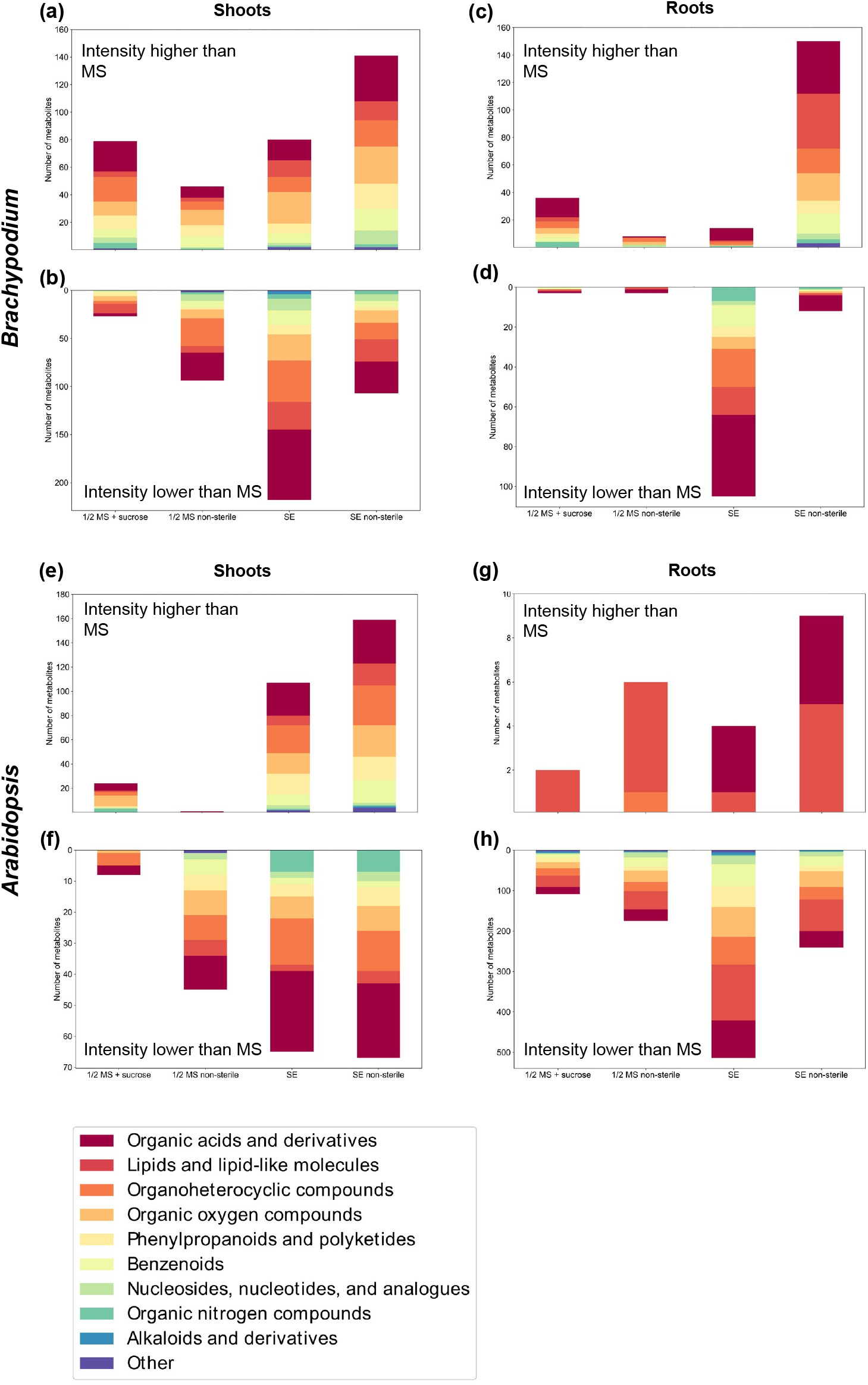
Higher or lower intensities of tissue metabolites for Brachypodium and Arabidopsis compared to ½ MS. (a-h) Number of metabolites changing significantly between ½ MS and other environments for Brachypodium (a-d) and Arabidopsis (e-h) tissues. Number of more and less abundant compounds are indicated on positive and negative axes, and metabolites are colored according to superclass. (Classyfire: https://cfb.fiehnlab.ucdavis.edu/, Djoumbou Feunang *et al*. (2016)).

**Figure S.5:**
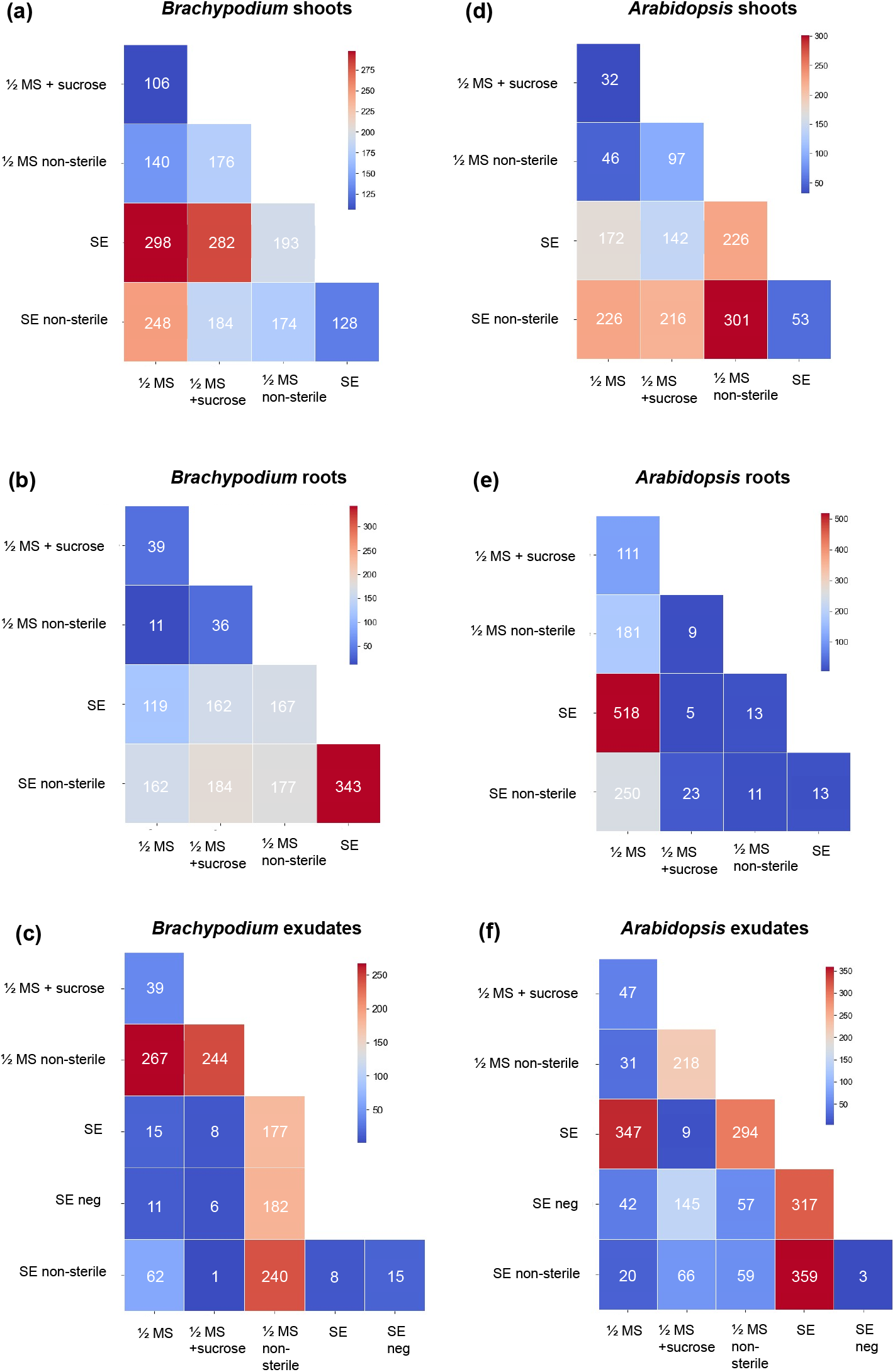
All pairwise comparisons of environmental conditions in Brachypodium and Arabidopsis. (a-c) Brachypodium pairwise comparisons. (d-f) Arabidopsis pairwise comparisons. Color code: red: high, blue: low.

**Figure S.6:**
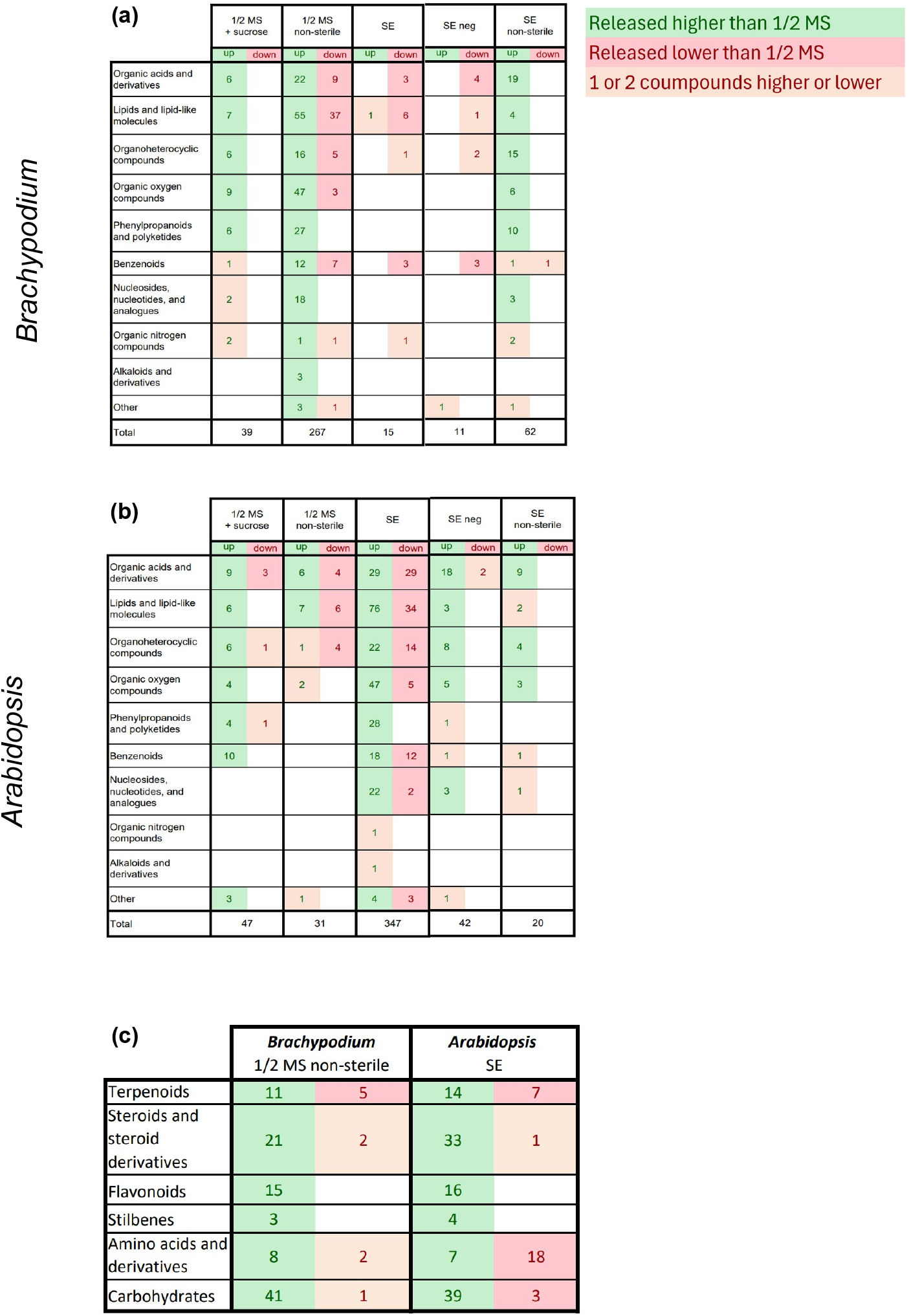
Elevated and diminished significant chemical classes in Brachypodium and Arabidopsis exudates in comparison to ½ MS. (a) Brachypodium showing up- and down-release patterns for all chemical classes in all environmental conditions. (b) Brachypodium showing up- and down-release patterns for all chemical classes in all environmental conditions. Color code: species release a higher quantity of a chemical class in comparison to ½ MS in green, species release a lower quantity in comparison to ½ MS in red. When 1 or 2 compounds of a chemical class are distinctly released, they are marked in orange.

**Figure S.7:**
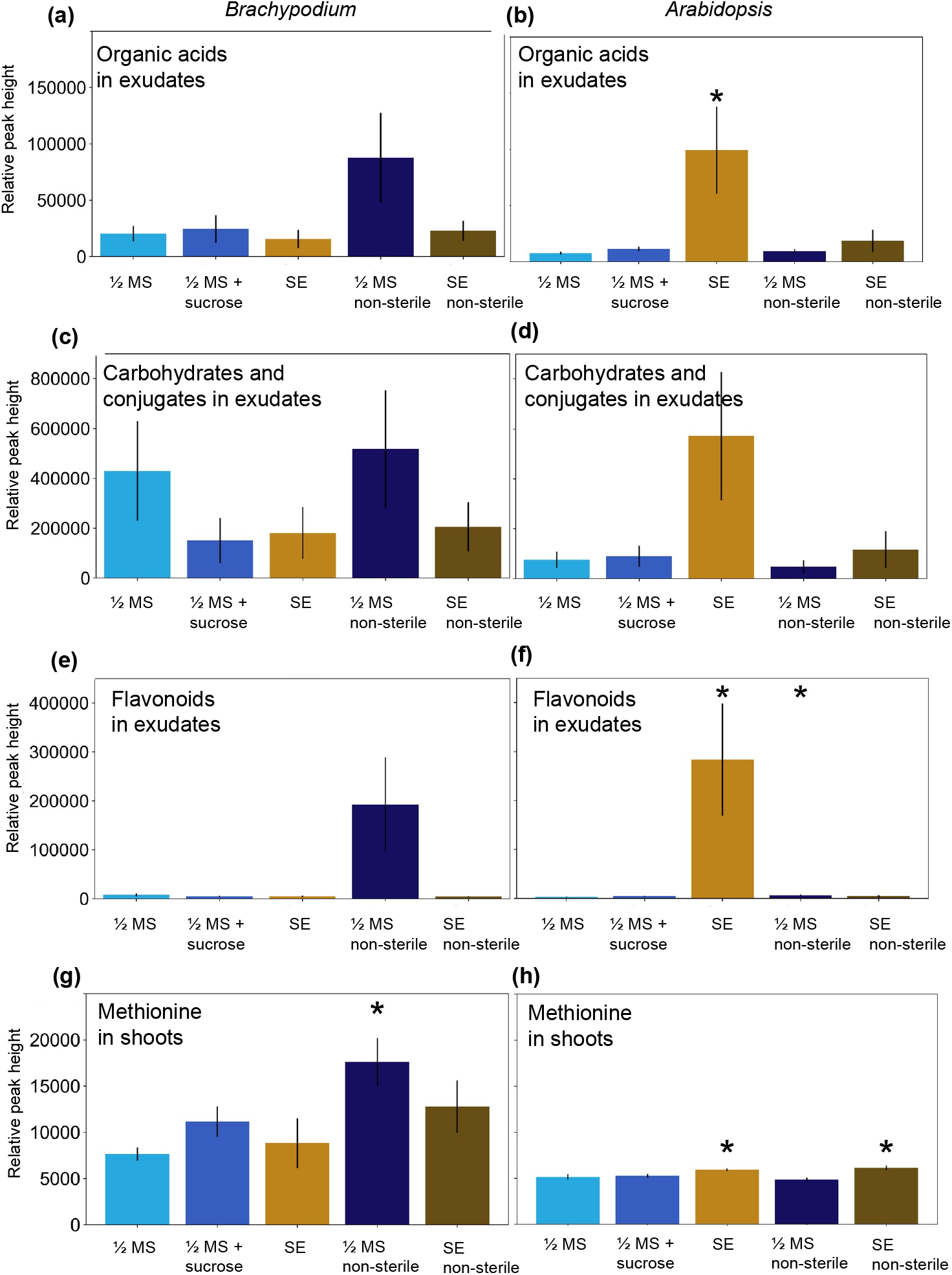
Primary and secondary metabolites accumulate in Brachypodium and Arabidopsis in ½ MS non-sterile and SE. (a-b) Organic acids in exudates of Brachypodium (a) and Arabidopsis (b). (c-d) Carbohydrates and conjugates in exudates of Brachypodium (c) and Arabidopsis (d). (e-f) Flavonoids in exudates of Brachypodium (e) and Arabidopsis (f). (g-h) Methionine in exudates of Brachypodium (g) and Arabidopsis (h). Data is displayed as average ± S.E.M., *: pvalue < 0.05 (T-Test vs. ½ MS).

